# The effects of sleep disruption, sex, and mating status on susceptibility to fungal (*Metarhizium anisopliae*) infections in *Drosophila melanogaster*

**DOI:** 10.1101/2025.07.21.665781

**Authors:** Mintong Nan, Raymond J. St. Leger

## Abstract

Important biological functions, including host defense, are linked to circadian rhythm and sleep. Studies have also indicated that the sex of the host can alter disease processes. In this study, we used a genetic approach in *Drosophila* to determine how sleep interacts with sex to influence the outcomes of infection with *Metarhizium anisopliae* strain Ma549. We found that as in mammals, *Drosophila* slept more after infection. The sleep-deprived *Drosophila Shaker* mutant with intact circadian rhythms failed to show this increased sleep response (sickness sleep) and succumbed quickly to infection. Mutants with disrupted PERIOD (PER) or CLOCK (CLK) affect circadian rhythms. The *per^01^*mutant shifted day/night cycles but not total sleep amounts, whereas the *Clk^Jrk^*mutant reduced sleep duration. Although mutations in *per* or *Clk* both impair sickness sleep, only *per* protects against disease, whereas active *Clk* reduces survival in infected *Drosophila,* which we attribute to it being pleiotropic. Independent of mutant status, males slept more than females, and virgins slept more than mated flies and survived infection longer, indicating that sex and mating status influence sleep and disease resistance irrespective of circadian rhythms.

## Introduction

Understanding the interplay between host genetic variation, physiological state, and pathogen resistance is fundamental to understanding the dynamics of host-pathogen interactions. The fruit fly *Drosophila melanogaster* is a widely used model organism for studying infection and host defense^1, 2^. Approximately 75% of genes responsible for human diseases have homologs in flies^3^. Many important disease-related genes in humans were first discovered in *Drosophila,* and the fly can also serve as a host for diverse human pathogens because of the remarkable conservation of pathogenesis and host defense mechanisms between insects and other animals^4^. Despite this conservation across phylogenetic distances, individual fruit flies and humans vary extensively from each other in their host responses to infectious diseases.

Fungi cause most insect diseases and infect insects by directly penetrating the cuticle using a combination of cuticle-degrading enzymes and mechanical pressure^5, 6^. However, host defenses against specialized entomopathogenic fungi are comparatively understudied. Most studies on fly immunity have tested bacterial or viral pathogens against fly mutants or deployed opportunistic mammalian pathogens injected directly into flies, bypassing several initial host defense barriers^4^. The entomopathogenic fungus used in this study, *Metarhizium anisopliae* ARSEF549 (Ma549), is naturally pathogenic to *Drosophila*^7–9^ and does not, like many other entomopathogen species, produce toxins that inhibit innate immune responses^10^. *Metarhizium* spp. have been used in host-defense studies to provide insights into emerging human pathogens^11–14^.

We previously demonstrated that approximately 9% of mutant *D. melanogaster* lines have altered disease resistance to Ma549^9^, and that DGRP (*Drosophila* Genetic Reference Panel made up of flies from originally isolated from a wild-caught population of *Drosophila melanogaster* collected in Raleigh, North Carolina, USA) flies differ greatly in longevity after Ma549 infection^8^. Our GWAS analysis of DGRP flies revealed a complicated network of immune and physiological genes, particularly sleep-related genes involved in *D. melanogaster* interactions with Ma549^8^. According to this GWAS analysis, a major predictor of how long a wild-caught North Carolinian fly will survive infections is its sleep pattern^8^. The worldwide variation in *Drosophila* susceptibility to disease will be a more complex phenomenon shaped by the interplay of host genetics, diet, environmental factors, and the specific types of pathogens locally present. However, we recently reported that resistance of a world-wide sampling of *Drosophila* lines to Ma549 is shaped by latitudinal gradients in climate (temperature, humidity) and sleep duration, with longer sleeping tropical males and virgin flies demonstrating increased survival during infection^15^.

These observations support the hypothesis that sleep and disease defense are intertwined traits linked to organismal fitness and subject to joint clinal evolution. Other studies in *Drosophila* have shown that mating reduces daytime but not nighttime sleep in uninfected females^16, 17^, and have documented sex-based differences or mating effects on immunity against various pathogens^8, 18–20^. Males of most *Drosophila* lineages are more resistant to Ma549 than are females, an effect mediated by immune responses^21^. However, the precise mechanisms underlying these effects, and their dependence on specific pathogens, remain unclear. Elucidating how host variation and sleep interact to influence immune defense against different pathogens, such as Ma549, is essential for developing a more holistic understanding of *Drosophila* immunity.

Sleep is an ancient behavior observed throughout the animal kingdom and is essential for survival in *Drosophila* as well as higher animals^22, 23^. Despite extensive research, the mechanism by which sleep enhances survival remains unclear. However, one hypothesis is that sleep deprivation has deleterious effects on immune functions^24^. Thus, the level of illness increases in night shift workers who report less sleep than daytime workers^25^. The dominant model to explain natural or healthy sleep that occurs with an approximately 24 h (circadian) periodicity is called the two-process model, whereby sleep duration is regulated by the interaction of two physiological processes: a circadian process tracking time of day and a homeostatic process tracking time awake and time asleep^26^. However, in addition to natural sleep, which occurs daily, several studies have identified sleep as a component of host defense that increases during an acute host response to infection^27^. This sickness sleep is regarded as a cardinal manifestation of sickness behavior, although the mechanisms and functions of sickness sleep are poorly understood^28^.

The conventional way to achieve sleep deprivation (SD) is by sensory stimulation^29^, and several mechanical devices that achieve SD with shaking have been applied to *Drosophila*^30^. However, the results have been unpredictable and non-reproducible, which may be attributable to damage to the flies and compensatory sleep increases^30–33^, which overestimate the role of sleep^29^. An alternative promising approach is genetic SD, which leads to more specifically targeted and robust sleep loss^29^. The development of genetic engineering tools has already led to the use of *Drosophila* models to investigate the role of circadian variation in the ability to fight bacteria^34–38^. Here, we utilized well-characterized sleep mutant lines to examine the role of sickness sleep and circadian clock-controlled genes in fruit fly responses to Ma549. The *period* (*per*) gene was first described in 1971^39, 40^, and along with the equally conserved transcription factor CLOCK, is critical for maintaining the circadian cycle in insects and vertebrates^39, 41^. In flies, CLK, along with another transcription factor CYCLE (CYC) forms the positive branch of the feedback loop, whereas PER and TIMELESS (TIM) form the negative branch^42^. The *Shaker* gene is involved in clock-independent control of sleep and facilitates plasticity of sleep in response to environmental conditions^43^. It encodes several voltage-gated potassium channel α-subunits and plays a key role in controlling neuronal membrane repolarization and transmitter release^44^.

In the current study, we demonstrate that infected flies increased daytime sleep soon after Ma549 infection, even before other overt signs of disease. Sleep mutants with an inactive *period* or *Shaker* gene had less sickness sleep and died significantly faster after infection, while a mutant with an impaired *Clock* gene showed enhanced survival. We explored the interaction of sleep/circadian genes with sex and mating status. Virgin flies of both sexes exhibited a lower fungal burden (within-host growth) than mated flies, which was surprising as any trade-offs produced by diverting energy and resources to egg production would not impact males. However, males had a lower fungal burden than females irrespective of mating and mutant status, indicating that sexual dimorphism in disease resistance is not causatively linked with differences in sleep patterns. These findings support our previous studies looking at wild fly populations^8, 15^ by highlighting the complex interplay between sleep, circadian genes, sex, mating, and fungal resistance in *Drosophila*.

## Results

### Ma549 infection promotes sickness sleep in the wildtype Canton-S line

We used *Drosophila* Activity Monitors (DAM2) following established protocols^45^ to analyze daytime and nighttime sleep in various fly lines, including Canton Special (CS) flies, a commonly used laboratory “wild-type” strain, and three sleep-affecting mutant strains, all of which were entrained to 12h:12h light:dark (LD) cycles^45^. The first 12 hours of sleep data were discarded to mitigate the effects of CO_2_ anesthesia and other acclimation effects. We monitored sleep-infection interactions from 12 to 60 hours post-infection, including two light-dark cycles. This period is before most flies showed overt symptoms such as reduced feeding and reproductive behavior, and before they up regulated the immune gene *Drosomycin* in response to Ma549 infection^9, 21^.

Consistent with a previous report^46^, mated Canton-S males and females had similar amounts of nighttime sleep (p = 0.7820, Fig. 1a-d and Supplementary Table S1-3a), but females slept 63.8% less than males during the daytime (ZT24-36 and ZT48-60, p < 0.0001, Fig. 1a-d and Supplementary Table S1-3a). Our initial investigation focused on determining whether infection alters sleep, with the aim of understanding the impact of infection on sleep metrics.

**Figure 1.**
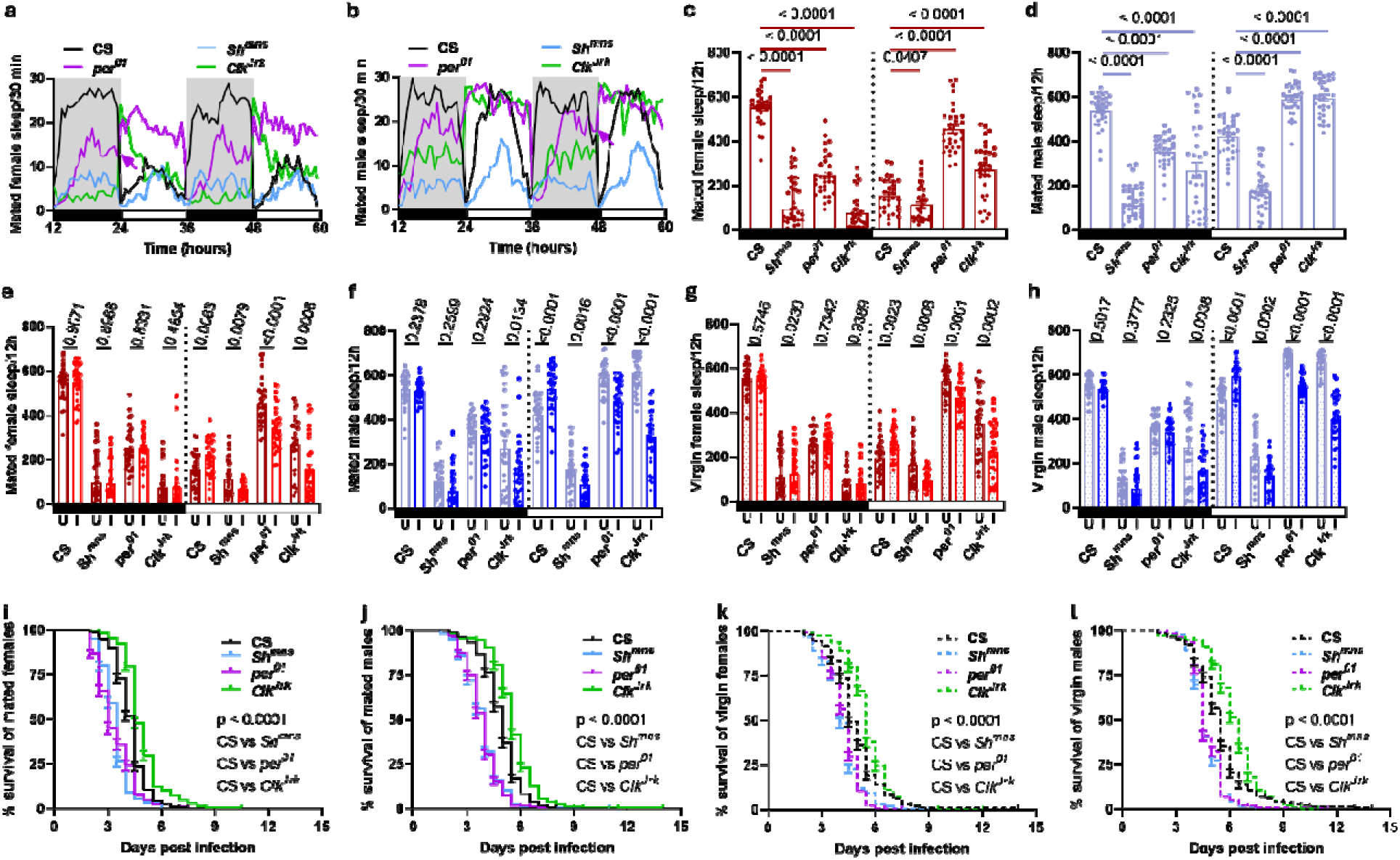
Ma549 infection affects sleep and survival. **Sleep profiles** show sleep in min per 30 min from 12-60 h for 32 flies pooled from three trials for uninfected mated CS (black), *Sh^mns^*(blue), *per^01^*(purple), *Clk^Jrk^*(green) females **(a)** and males **(b)**. Gray shading indicates nighttime. **Sleep comparisons included uninfected mated CS versus mutant females (c) or males (d), as well as uninfected (U) versus infected (I) mated females (e), mated males (f), virgin females (g), and virgin males (h) across the CS/*Sh^mns^*/*per^01^*/*Clk^Jrk^* lines.** Bars depict nighttime (black) and daytime (white) sleep for uninfected (dark red for females, sky blue for males) and infected (scarlet red for females, navy blue for males) mated and virgin (shaded) flies pooled from three trials. Each data point represents the average sleep/12h of a single fly (n = 32) from 12-60 h post infection. In panels **c-d**, statistical analyses included the Mann-Whitney U test for nighttime sleep in *per^01^* males, Welch’s t-tests for nighttime sleep in *Sh^mns^* females and *Clk^Jrk^* males, and for daytime sleep in *Clk^Jrk^* females. Unpaired Student’s t-tests for all other comparisons. In panels **e-h**, Mann-Whitney U tests for nighttime sleep in mated CS, *Sh^mns^* flies, virgin *Sh^mns^*females, and *Clk^Jrk^* females, and Welch’s t-tests for nighttime sleep in virgin CS males, *Clk^Jrk^* males, and for daytime sleep in mated *Sh^mns^* females, virgin *Sh^mns^*flies, and virgin *Clk^Jrk^* males. Unpaired Student’s t-tests for all other comparisons. The mean ± SEM and p-values are indicated. **Differences in survival after Ma549 infection.** Kaplan-Meier plots illustrate survival disparities between CS (black) and *Sh^mns^*(blue), or *per^01^*(purple), or *Clk^Jrk^* (green) in mated females (n = 230-321) (**i**), mated males (n = 268-329) (**j**), virgin females (n = 288-346) (**k**), and virgin males (n = 281-345) (**l**). Statistical significance was determined using the log-rank and Wilcoxon tests for intergroup comparisons.

Compared to uninfected CS controls, Ma549 infection significantly increased daytime sleep in mated or virgin females and males between 21.6% and 36.4% (p ≤ 0.0063, Fig. 1e-h and Supplementary Table S2, 3c). Following precedent in other animals, including humans, we refer to this increase in sleep following infection as sickness sleep. However, Ma549 infection did not significantly alter nighttime sleep in either sex, regardless of mating status (p ≥ 0.2378, Fig. 1e-h and Supplementary Table S2, 3c). These findings indicate that Ma549 infection promotes daytime sickness sleep in the CS line.

### Sickness sleep is disrupted in short-sleeping *Shaker* line and clock gene mutants, with variation in survival to Ma549 infection

We investigated three well-characterized mutant fly lines: *Sh^mns^*, *per^01^*, and *Clk^Jrk^*, carrying inactivated *Shaker (Sh)*, *period (per)*, and *Clock (Clk)* genes, respectively. Mated mutant flies were compared with wild-type CS controls to establish and verify baseline sleep profiles under normal conditions. When analyzing the sleep of infected mutants, we compared them with their uninfected peers to accurately assess the impact of infection on sleep within each mutant line. We examined male and female sickness sleep and survival after Ma549 infection to examine potential sex-specific differences in sleep, immune response, and disease outcomes. Additionally, we compared mated and virgin flies to determine whether mating status alters disease resistance and sleep. To quantify the sleep difference between groups within the same fly line, we calculated the net percentage change in sleep duration of group B relative to group A. This involved subtracting group A sleep from group B sleep and then normalizing by dividing by the group A value (Supplementary Table S3).

*Sh^mns^* flies are characterized by a *Shaker* gene mutation that affects the amount of sleep but not circadian regulation^44^. Compared with uninfected CS flies, mated *Sh^mns^*females (males) slept 74.5% (77.2%) less at night and 26.6% (59.1%) less during the day (p ≤ 0.0407, Fig. 1a-d and Supplementary Table S1, 2, 3b). This resulted in a significant two-thirds reduction in total sleep compared to wild-type CS flies, consistent with a previous report^44^. Compared to uninfected *Sh^mns^*flies, Ma549 infection significantly reduced daytime sleep in mated and virgin *Sh^mns^* females and males by between 38.0% and 40.4% (p ≤ 0.0079, Fig. 1e-h and Supplementary Table S2, 3c). However, Ma549 infection did not significantly alter nighttime sleep in either sex, regardless of mating status (p ≥ 0.2599, Fig. 1e-h and Supplementary Table S2, 3c). Therefore, while Ma549 infection promoted daytime sleep of the CS line, both mated and virgin *Sh^mns^*flies showed reduced daytime sleep post-infection compared to their uninfected peers, suggesting a heightened susceptibility to sleep disruption.

Given the central role of CLOCK and PERIOD in regulating circadian rhythms, using *per^01^* and *Clk^Jrk^* mutants provides a unique opportunity to further study the interaction between Ma549 infection and sleep. *Per^01^* flies harbor a stop codon in *period* that disrupts the circadian cycle in both sexes^40^ so they exhibit fluctuating sleep/woke behavior under light:dark conditions but display complete arrhythmicity in constant darkness^47^. Our results showed that uninfected mated *per^01^*females (males) had a significant decrease of 55.5% (34.8%) in nighttime sleep and a significant increase of 198.1% (39.7%) in daytime sleep compared to CS flies (p < 0.0001, Fig. 1a-d and Supplementary Table S1, 2, 3b). Reduced nighttime and increased daytime sleep evened out, such that total sleep per 24 h was not significantly reduced in the *per^01^* mutants of either sex. Similar to previous reports^40, 47^, we found that *per^01^* flies exhibited fluctuating sleep patterns because of the loss of the light-off to light-on transition (arrows shown in Fig. 1a-b), albeit with periodic behaviors in LD cycles. Compared to uninfected *per^01^* flies, Ma549 infection did not significantly alter nighttime sleep of *per^01^* (p ≥ 0.2325, Fig. 1e-h and Supplementary Table S2, 3c). However, it reduced daytime sleep by between 14.1% and 25.5% in mated and virgin females and males (p ≤ 0.0001, Fig. 1e-h and Supplementary Table S2, 3c). Thus, following Ma549 infection, the nighttime sleep of the *per^01^* mutants remained unaffected, while daytime sleep was significantly reduced compared to that of uninfected peers, like the pattern observed in *Sh^mns^* flies.

The *Clk^Jrk^* arrhythmic mutant carries a premature stop codon that eliminates much of the activation domain of this transcription factor, and it only expresses low levels of PERIOD and TIMELESS^48^. Resembling a previous report^48^, we found that *Clk^Jrk^* flies did not react to the lights-on transition (Fig. 1a-b). Compared to uninfected CS flies, uninfected *Clk^Jrk^* females (males) showed an 86.2% (50.3%) reduction in nighttime sleep, but a 77.7% (41.6%) increase in daytime sleep (p < 0.0001, Fig. 1a-d and Supplementary Table S1, 2, 3b). Thus, *Clk^Jrk^* mutants sleep much less at night and much more during the day, indicating a shift in the day-night sleep cycle. Compared to uninfected *Clk^Jrk^* flies, daytime sleep decreased following infection between 36.6% and 46.8% in mated or virgin females and males (p ≤ 0.0006, Fig. 1e-h and Supplementary Table S2, 3c). However, nighttime sleep remained unaffected in females (p ≥ 0.4854, Fig. 1e-h and Supplementary Table S2, 3c) but showed a significant reduction of approximately 42% in both mated and virgin males (p ≥ 0.0134, Fig. 1e-h and Supplementary Table S2, 3c), suggesting a sex-specific effect of disrupting the day-night cycle. Thus, while Ma549 infection primarily affects daytime sleep in *Clk^Jrk^* mutants, nighttime sleep is significantly reduced only in males.

Longevity following Ma549 infection varied significantly between the mutant flies and wild-type CS. Compared to infected CS flies, mated (virgin) *Sh^mns^* flies of both sexes died approximately 21.4% (16.3%) faster, and similarly *per^01^* flies died approximately 22.3% (14.5%) faster (p < 0.0001, Fig. 1i-l and Supplementary Table S4), showing that active *Shaker* and *per* genes both protect against fungal infection. In contrast, mated (virgin) *Clk^Jrk^* flies of both sexes outlived wild-type CS flies by approximately 15.8% (13.0%) (p < 0.0001, Fig. 1i-l and Supplementary Table S4), indicating that a functioning *Clk* gene reduces longevity.

### Sexual dimorphism in sleep and longevity in response to Ma549 infection

Next, we specifically examined the effects of sexual dimorphism on sleep and longevity after Ma549 infection. Only males with clock-disrupted *per^01^* and *Clk^Jrk^* mutations exhibited significantly (p ≤ 0.0008) more nighttime sleep following infection, increasing their overall sleep compared to that of infected females (Fig. 2a & f and Supplementary Table S2, 3d). Neither mated nor virgin CS or *Sh^mns^* males differed significantly (p ≥ 0.1562) from females in nighttime sleep (Fig. 2a & f and Supplementary Table S2, 3d). However, daytime sickness sleep varied according to sex in all lines. Specifically, compared to infected mated (virgin) females, infection increased daytime sleep in mated (virgin) males by 158.8% (130.4%) in CS, 49.4% (46.3%) in *Sh^mns^*, 40.9% (18.5%) in *per^01^*, and 105.1% (78.6%) in *Clk^Jrk^* (p ≤ 0.0198, Fig. 2a & f and Supplementary Table S2, 3d). Thus, as previously stated, Ma549 infection reduced daytime sleep in mutant strains but enhanced it in the CS line (Fig. 1e-h and Supplementary Table S2, 3c). However, these results also show that infected male flies sleep significantly more than female flies, highlighting sex-specific differences in sleep responses to infection and variations across different genetic strains.

**Figure 2.**
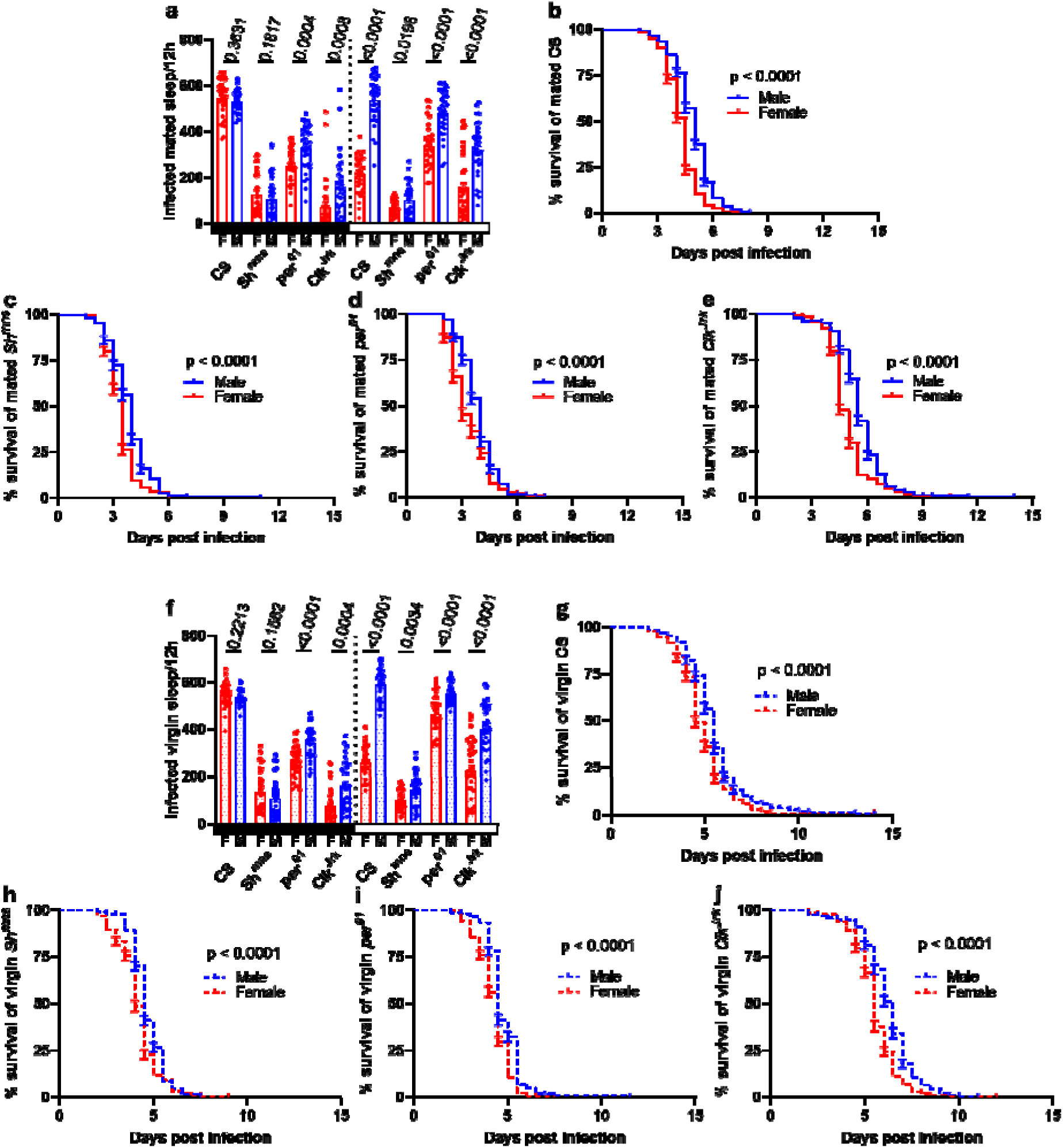
Sexual dimorphism in sleep and longevity against Ma549 infection in CS, *Sh^mns^*, *per^01^*, and *Clk^Jrk^* flies. Comparison of sickness sleep between females and males in mated (a) and virgin (f) groups. Each data point represents the average sleep/12 h of a single fly (n = 32 flies) during nighttime and daytime from 12-60 h post infection pooled from three trials. Statistical significance was assessed for comparisons between the female and male groups within CS, *Sh^mns^*, *per^01^*, and *Clk^Jrk^* lines. For nighttime sleep, Welch’s t-test was used for mated and virgin CS, and virgin *Clk^Jrk^*, with the Mann-Whitney U test for mated *Sh^mns^* and *Clk^Jrk^*. For daytime sleep, Welch’s t-test was used for mated and virgin *Sh^mns^*, and virgin *per^01^*. The unpaired Student’s t-test was used for all other comparison groups. The mean ± SEM and p-values are indicated. Black (white) rectangles on the x-axis indicate nighttime (daytime), whereas shaded bars denote virgin flies. Line Abbreviations: F for females and M for males. **Survival differences between males and females after Ma549 infection.** Kaplan-Meier plots illustrate the survival disparities between females (n = 230-346) and males (n = 268-345) of mated CS (**b**), *Sh^mns^* (**c**), *per^01^* (**d**), *Clk^Jrk^*(**e**), and virgin CS (**g**), *Sh^mns^* (**h**), *per^01^*(**i**), *Clk^Jrk^* (**j**). Statistical significance at p < 0.0001 was determined using both the log-rank test and Wilcoxon test for comparisons between two groups.

Interestingly, differences in sleep patterns between males and females translated into differences in longevity. Mated (virgin) males in the CS, *Sh^mns^ per^01^*, and *Clk^Jrk^* lines outlived mated (virgin) females by 15.0% (13.9%), 15.8% (12.7%), 16.6% (13.2%), and 14.3% (12.6%), respectively (p < 0.0001, Fig. 2b-e, Fig. 2g-j, and Supplementary Table S4).

Taken together, our findings suggest that males, irrespective of mating status and mutant line, exhibit longer sickness sleep and greater disease resistance than females following Ma549 infection.

### Impact of mating status on sleep and survival in flies following Ma549 infection

Next, we analyzed the data with the goal of disentangling the effect of mating status on fly sleep and survival following Ma549 infection. Mating status did not affect the nighttime sleep of infected CS, *Sh^mns^*, *per^01^*, and *Clk^Jrk^* females or males (p ≥ 0.1626, Fig. 3a & f and Supplementary Table S2, 3e). However, mating significantly reduced daytime sleep across all strains irrespective of sex and infection status. Specifically, infection reduced daytime sleep in mated females (males) by 19.2% (9.2%) in the CS line, 29.5% (28.0%) in *Sh^mns^*, 27.5% (13.8%) in *per^01^*, and 30.8% (20.6%) in *Clk^Jrk^* (p ≤ 0.0354, Fig. 3a & f and Supplementary Table S2, 3e).

**Figure 3.**
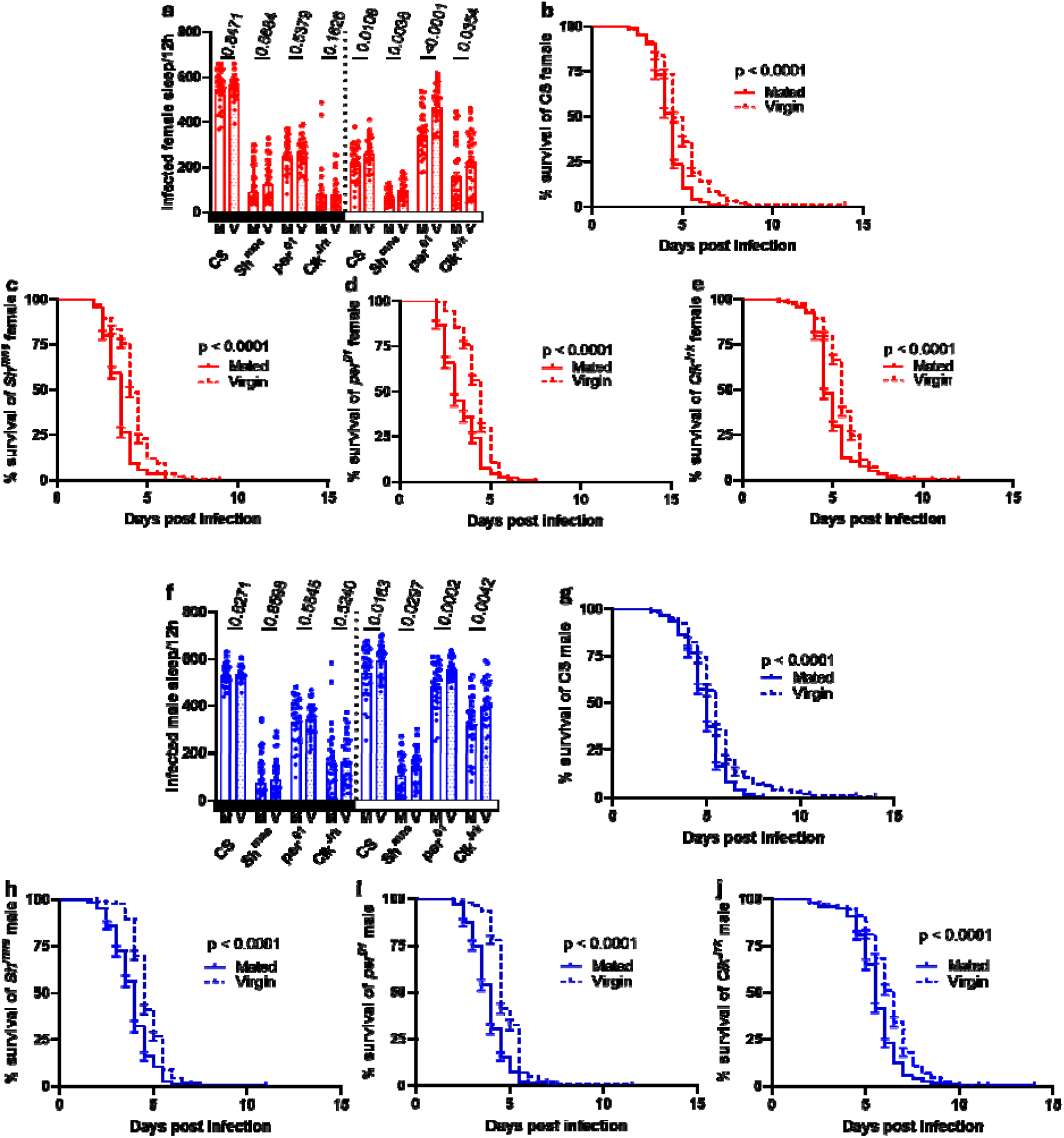
Mating status affects sleep and longevity following Ma549 infection in CS, *Sh^mns^*, *per^01^*, and *Clk^Jrk^* flies. Differences in nighttime and daytime sleep between mated and virgin flies in the infected female (a) and male (f) groups. Each data point represents the average sleep per 12 h of a single fly (n = 32 flies) during the nighttime and daytime from 12-60 h post infection. Statistical significance was assessed for comparisons between mated and virgin females or males within the CS, *Sh^mns^*, *per^01^*, and *Clk^Jrk^* fly lines. For nighttime sleep, the unpaired Student’s t-test was used for infected *per^01^* females and males, with Welch’s t-test for infected CS females and males. The Mann-Whitney U test was used for all other comparison groups. For daytime sleep, Welch’s t-test was used for infected CS and *per^01^* males, and the unpaired Student’s t-test was used for all other comparison groups. The mean ± SEM and p-values are indicated. Black (white) rectangles on the x-axis indicate nighttime (daytime), whereas shaded bars denote virgin flies. Line Abbreviations: V for virgin and M for mated. **Survival differences between mated and virgin flies after Ma549 infection.** Kaplan-Meier plots illustrate the survival disparities between mated (n = 230-329) and virgin (n = 281-346) flies of CS (**b**), *Sh^mns^*(**c**), *per^01^* (**d**), and *Clk^Jrk^* females (**e**), as well as CS (**g**), *Sh^mns^* (**h**), *per^01^*(**i**), and *Clk^Jrk^*(**j**) males. Statistical significance at p < 0.0001 was determined using both the log-rank test and Wilcoxon test for comparisons between two groups.

After infection, virgin females (males) outlived their mated peers by 16.7% (15.6%) in CS, 25.4% (22.0%) in *Sh^mns^*, 29.7% (25.9%) in *per^01^*, and 14.2% (12.5%) in *Clk^Jrk^* (p < 0.0001, Fig. 3b-e & 3g-j and Supplementary Table S4). These findings reveal a notable impact of mating status on the regulation of sleep and immunity in flies, although these effects are independent of mutant status.

### Effects of line, sex, mating status, and infection on sleep

To quantify the effect of infection on sleep across various fly genotypes, we calculated the net percentage change in sleep duration of infected flies compared to controls. This involved subtracting uninfected control sleep from infected sleep and then normalizing by dividing by the control value from the same fly line (Fig. 4a-b). This approach accounts for inherent sleep variations between fly lines and potential disruptions in circadian rhythms caused by mutations. Consequently, it offers a more refined and statistically robust method for comparing the effects of infection on sleep across different genotypes; it accounts for inherent sleep differences across genotypes and helps isolate infection-driven effects from baseline variability.

**Figure 4.**
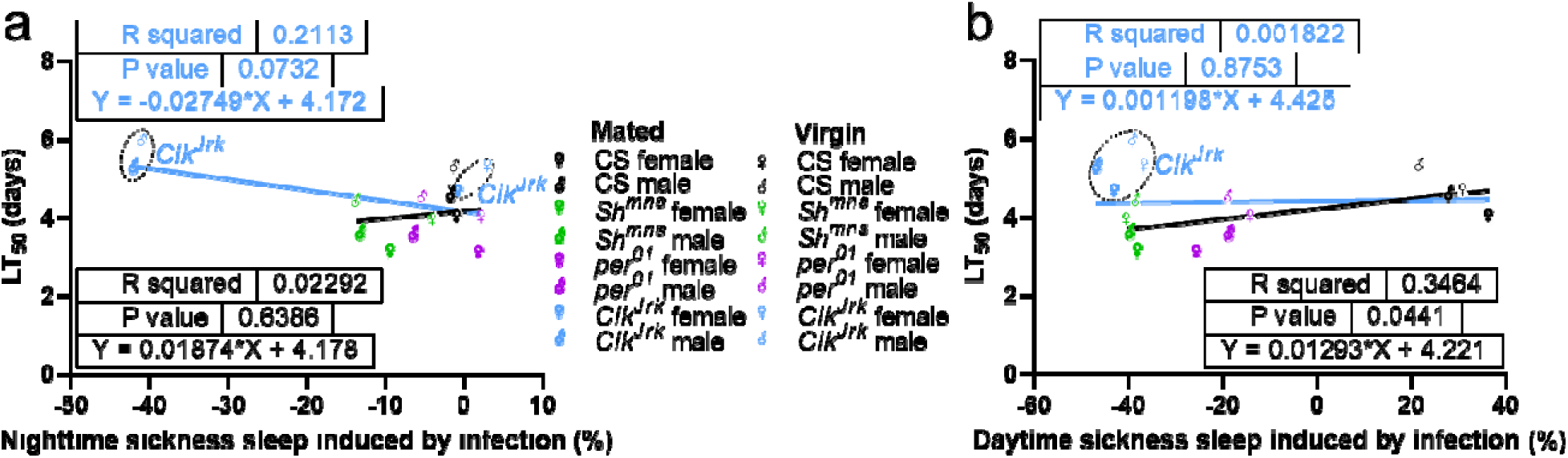
Correlations between LT_50_ values and net percentage change in nighttime (a) or daytime (b) sleep duration of infected flies compared to controls were analyzed for mated and virgin CS (black symbols), *Sh^mns^*(green symbols), *per^01^* (purple symbols), and *Clk^Jrk^*(blue symbols) females (♀) and males (♂). The light blue regression line includes *Clk^Jrk^* (circled blue symbols), while the black regression lines do not. The figures display R^2^ values, p-values, and regression equations with slopes.

We observed no correlation between the infection-induced net percentage change in nighttime or daytime sleep and disease resistance when all four groups—mated and virgin males and females across the four mutant lines—were included in the regression (blue lines; Fig. 4a–b). However, to avoid overgeneralizing by combining all mutants in a single analysis, we accounted for their distinct differences in sleep architecture and circadian function. *Per^01^* represents the negative arm of the circadian clock, *Clk^Jrk^* is part of the positive arm, and so non-circadian effects, if any, may be in opposite directions. We excluded *Clk^Jrk^*from follow-up analyses to prevent skewed interpretations driven by this specific genotype. These refined analyses revealed a strong positive correlation between infection-induced net percentage change in daytime sleep and disease resistance—but not in nighttime sleep (black lines; Fig. 4a–b)—suggesting that infection-induced changes in daytime sleep and disease outcomes are not independently regulated.

Next, we used analysis of variance (ANOVA) to investigate the interplay of genetic line, sex, mating status, and infection on nighttime (Fig. 5a) and daytime (Fig. 5b) sleep (Table 1 generated from data in Supplementary Table S2). Both night and daytime sleep patterns exhibited highly significant main effects of genetic line (p < 0.0001, Table 1 and Fig. 5), confirming that CS, *Sh^mns^*, and *per^01^*showed distinct sleep behaviors. Males and females differed significantly (p ≤ 0.0006) in both night and daytime sleep durations (Table 1 and Fig. 5), consistent with sexual dimorphism in sleep patterns. Infection and mating status differentially affected sleep patterns, with both factors significantly reducing daytime sleep (p < 0.0001), while their effects on nighttime sleep were less pronounced. Infection had a marginal effect on nighttime sleep (p = 0.0878), and mating status had no significant impact during the night (p = 0.2777), highlighting a time-of-day specificity in how these biological factors influence sleep (Table 1 and Fig. 5). Daytime sleep exhibited significant interactions between line and each of sex, mating status, and infection (all p ≤ 0.0005). In contrast, nighttime sleep showed a significant interaction only between line and sex (p < 0.0001; Table 1 and Fig. 5). Additionally, a significant three-way interaction among line, sex, and infection was detected for daytime sleep (p = 0.0066), but not for nighttime sleep (p = 0.6316), suggesting that the influence of any one factor on daytime sleep can be substantially modified by the presence of the others (Table 1 and Fig. 5).

**Figure 5.**
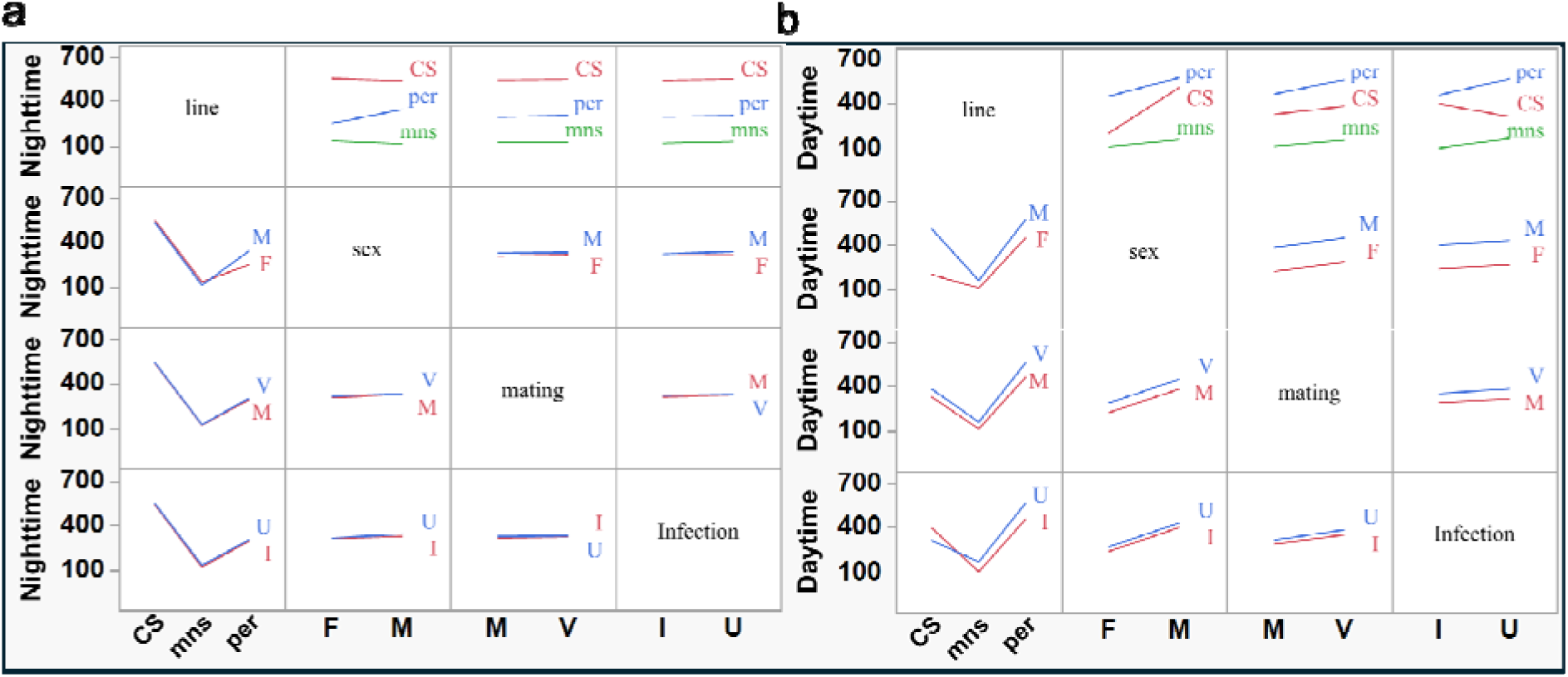
Interaction profiles of line, sex, mating status, and infection status on nighttime (a) and daytime (b) sleep across CS, *Sh^mns^*, and *per^01^*females and males. The interaction profiles of line, sex, mating status, and infection status on nighttime and daytime sleep (minutes/12hr) across the three genetic lines (CS, *Sh^mns^*, and *per^01^*), both sexes (males and females), two mating statuses (mated and virgin), and two infection statuses (infected and uninfected) were generated from the sleep data of 768 individual flies. Abbreviations: sex (F = female, M = male), mating status (M = mated, V = virgin), and infection status (I = infected, U = uninfected).

**Table 1:** Effects of line, sex, mating status, and infection as well as their interactions on night and daytime sleep. (Source: This is a list of the tested factors and interactions. Nparm: Number of parameters associated with the effect. DF: degrees of freedom for each source of variation. Sum of Squares: sum of squares (SS) for each source of variation, along with the total from all sources. F Ratio: the mean square of the factor (lot) divided by the mean square of the error. A p-value less than 0.05 means that the factor or interaction has a statistically significant effect.)

| Source | Nparm | DF | Nighttime |  |  | Daytime |  |  |
| --- | --- | --- | --- | --- | --- | --- | --- | --- |
|  |  |  | Sum of squares | F ratio | p values | Sum of squares | F ratio | p values |
| line | 2 | 2 | 22488250 | 1916.522 | <0.0001 | 18176551 | 1448.466 | <0.0001 |
| sex | 1 | 1 | 70553 | 12.0254 | 0.0006 | 4951748 | 789.1969 | <0.0001 |
| mating | 1 | 1 | 6924 | 1.1802 | 0.2777 | 822942 | 131.1584 | <0.0001 |
| Infection | 1 | 1 | 17148 | 2.9228 | 0.0878 | 173010 | 27.5739 | <0.0001 |
| line*sex | 2 | 2 | 526654 | 44.8831 | <0.0001 | 2185484 | 174.1584 | <0.0001 |
| line*mating | 2 | 2 | 2097 | 0.1787 | 0.8364 | 96281 | 7.6725 | 0.0005 |
| line*Infection | 2 | 2 | 1250 | 0.1066 | 0.8989 | 1315491 | 104.8298 | <0.0001 |
| sex*mating | 1 | 1 | 207 | 0.0353 | 0.8510 | 3.12630208 | 0.0005 | 0.9822 |
| sex*Infection | 1 | 1 | 6487 | 1.1056 | 0.2934 | 151 | 0.0240 | 0.8770 |
| mating*Infection | 1 | 1 | 129 | 0.0220 | 0.8821 | 1371 | 0.2185 | 0.6404 |
| line*sex*mating | 2 | 2 | 462 | 0.0394 | 0.9614 | 13309 | 1.0605 | 0.3468 |
| line*sex*Infection | 2 | 2 | 5396 | 0.4598 | 0.6316 | 63457 | 5.0568 | 0.0066 |
| line*mating*Infection | 2 | 2 | 136 | 0.0116 | 0.9885 | 8903 | 0.7094 | 0.4923 |
| sex*mating*Infection | 1 | 1 | 0.52083333 | 0.0001 | 0.9925 | 6966 | 1.1102 | 0.2924 |
| line*sex*mating*Infection | 2 | 2 | 395 | 0.0337 | 0.9669 | 7428 | 0.5920 | 0.5535 |

### The impact of natural host variation and sleep mutants on fungal fitness

To investigate the impact of host variation on fungal fitness, we compared within-host growth [fungal load, measured as colony forming units (CFU)] between the CS and sleep mutant lines. The time course of CFU counts showed that irrespective of mating status, CS and *Clk^Jrk^* flies delayed fungal growth compared to the more susceptible *Sh^mns^* and *per^01^* flies, although CS flies exhibited higher fungal loads than the more resistant *Clk^Jrk^* flies (Fig. 6a-c generated from data in Supplementary Table S5). CFUs/fly ranged from ≤ 8.4 ± 1.5 in virgin male *Clk^Jrk^*flies at day 4.5 to ≥ 9149.0 ± 244.7 in the susceptible lines at day 2.5, showing that the susceptible sleep mutant lines, particularly if they were female and mated, were much less able to restrain fungal growth. CFUs appeared 1.5 days post-infection in the particularly susceptible mated *per^01^*and *Sh^mns^* females, with an average of 40.5 ± 5.6 and 38.2 ± 5.2 CFUs/fly, respectively. In contrast, CFUs appeared 2.5 days post-infection in wild-type CS females, but remained at low levels (< 5 per fly) until day 3. CFUs appeared 4.5 days post infection in the most resistant virgin *Clk^Jrk^* males, but remained ≤ 20 per fly until day 5. In all four lines, fungal loads only climbed rapidly when flies were close to death, indicative of host resistance breaking down.

**Figure 6.**
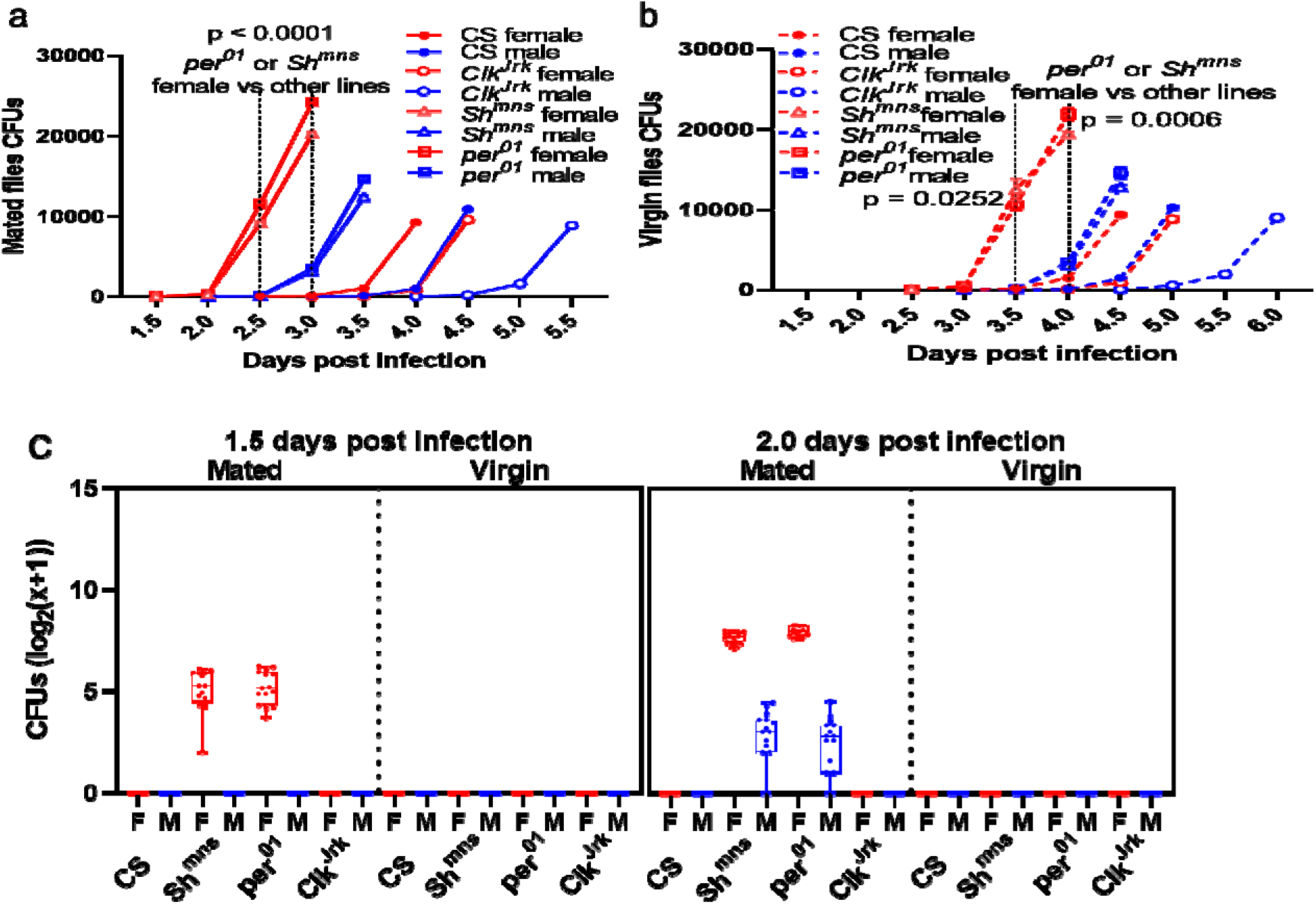
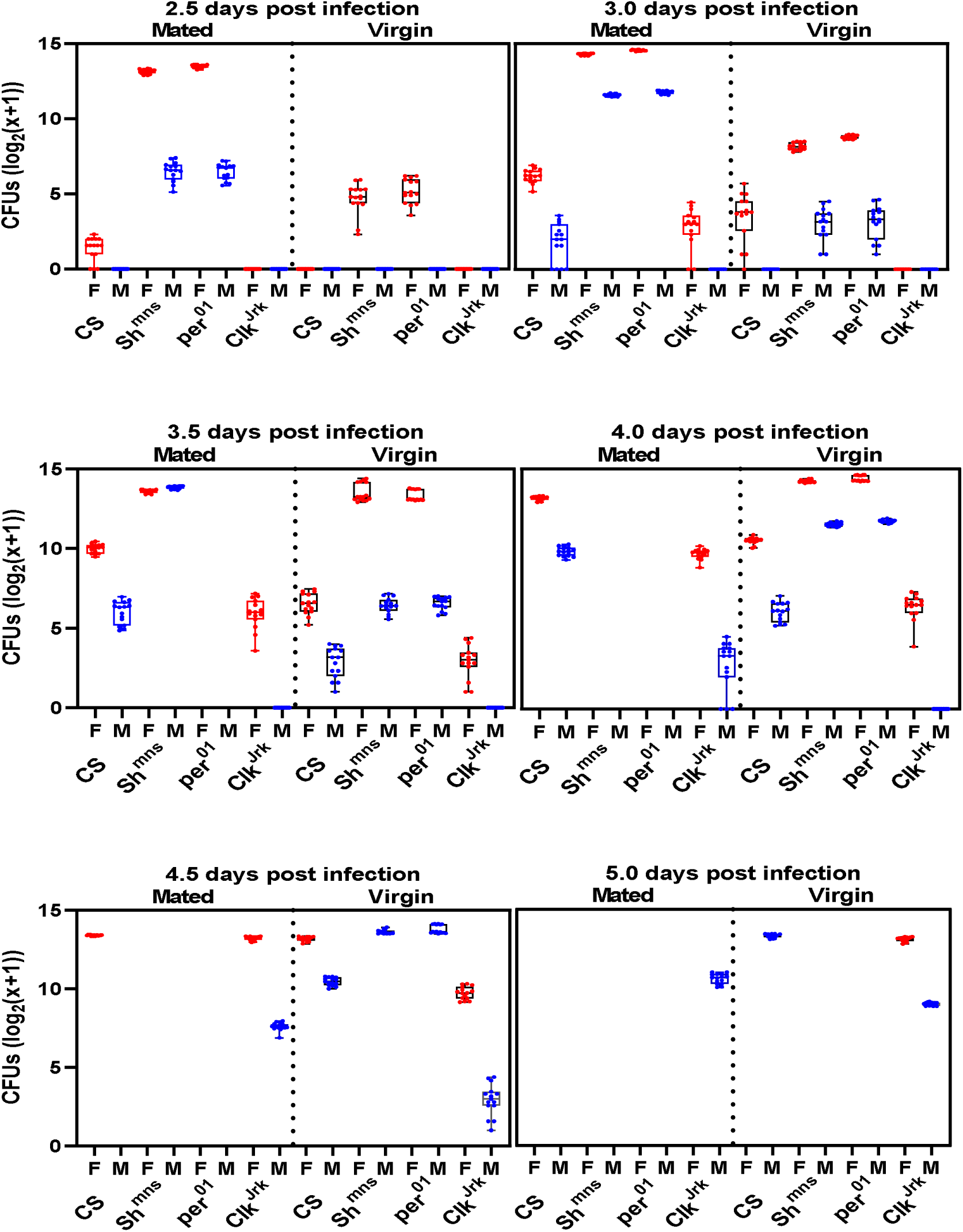
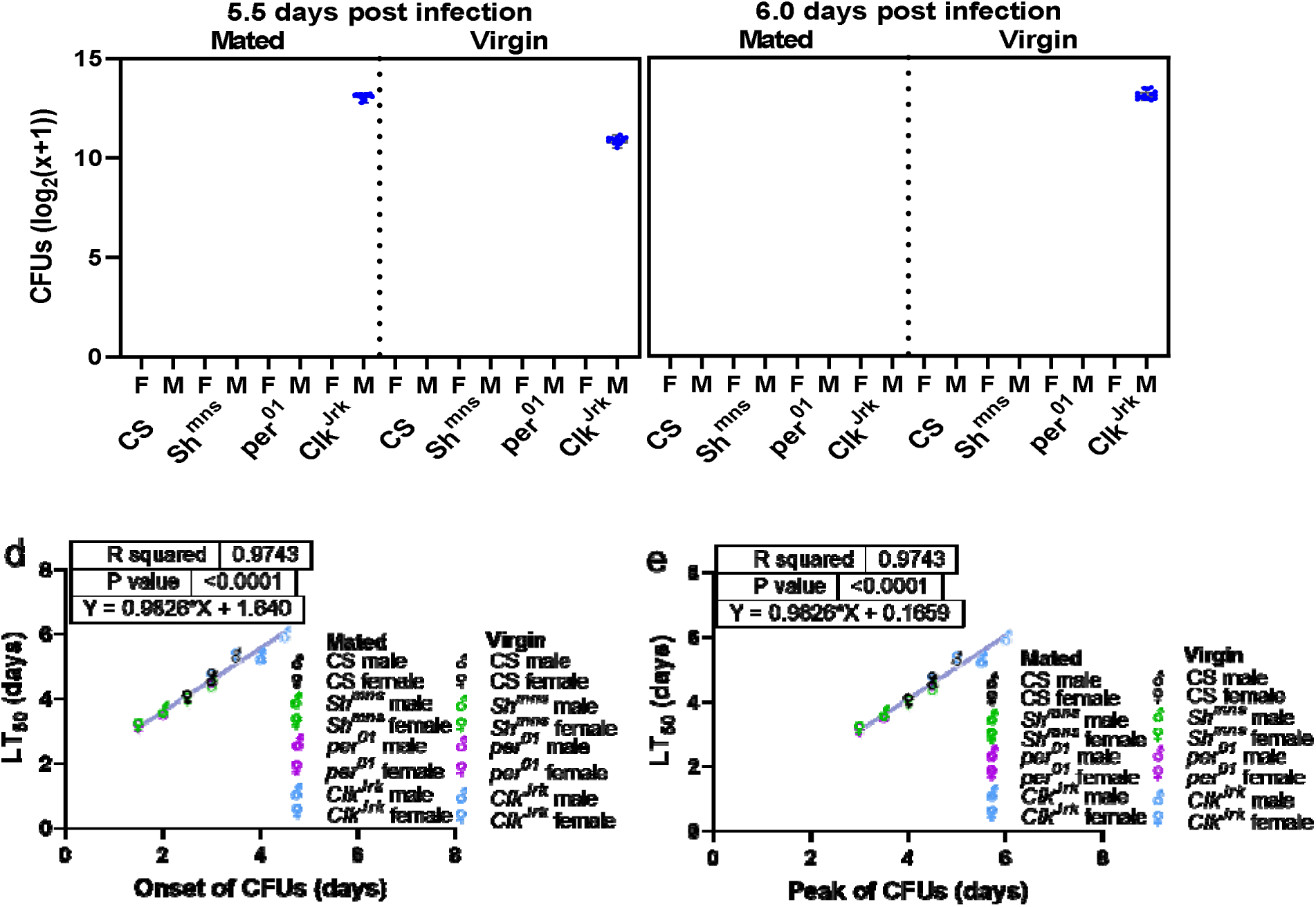
Time course of CFU (colony forming units) production in mated (a) and virgin (b) CS and mutant flies following Ma549 infection. Flies were individually homogenized and plated at 12-hour intervals until death. Each data point represents the average of CFU values of one of the CS, *Sh^mns^*, *per^01^*, and *Clk^Jrk^* lines from 15 flies, with five replicates each, per sex, mating status, line, and time point across three experiments. Means ± SEM and p-values were obtained from two-way ANOVA and are indicated. **Time course of log-transformed Ma549 fungal loads in the hemolymph of sleep mutants and their corresponding isogenic WT backgrounds (c).** Results are depicted as box plots for each time point, with the mean ± SEM of CFUs presented for 15 individual replicates per sex, mating status, line, and time point across three repeated experiments. Correlations between LT_50_ values and CFU onset (d) or peak (e) were analyzed for infected mated and virgin CS (black symbols), *Sh^mns^* (bright green symbols), *per^01^*(electric purple symbols), and *Clk^Jrk^* (dodger blue symbols) females (♀**) and males (**♂). The figures display R^2^ values, p-values, and regression equations with slopes.

Irrespective of mating status, males exhibited a lower fungal load than females, with proliferation delayed approximately 0.5-1 day (Fig. 6a-c), aligned with LT_50_ values (Fig. 2b-e, g-j). Likewise, proliferation was delayed in virgin flies by approximately 0.5-1 day relative to mated flies (Fig. 6b, c), also aligned with LT_50_ values (Fig. 3b-e, g-j). ANOVA revealed significant differences in fungal loads on days 2.5 and 3.0 between mated *per^01^* or *Sh^mns^* females and mated females from other lines, as well as all males across the four lines studied (p < 0.0001 Fig. 6a). Additionally, significant differences were found on days 3.5 and 4.0 between virgin *per^01^* or *Sh^mns^* females and virgin females from other lines, along with all males across the four lines studied (p ≤ 0.0252, Fig. 6b). In addition, regression analyses demonstrated that the LT_50_ values (Supplementary Table S4b) positively correlated with both the onset (initial time point at which fungal growth becomes detectable) and peak time (time point maximum fungal growth achieved) of CFUs (R² = 0.9743, p < 0.0001, Fig. 6d-e) across mated and virgin CS, *Sh^mns^*, *per^01^*, and *Clk^Jrk^*females and males. Thus, flies with delayed onset or peak times of fungal load survived longer.

These results indicate that the fungal load in the hemocoel is affected by the fly’s genetically determined ability to regulate sleep, and proliferation is coincident with and presumably causes death; that is, the flies cannot tolerate high levels of fungi (Fig. 6).

## Discussion

Infection, mating, and sex are critical factors that intricately influence sleep and survival in various species. Infection often triggers an immune response that can significantly alter sleep patterns so as to conserve energy and prioritize the body’s resources toward fighting off pathogens^28^. This adaptive change in sleep architecture serves as a vital mechanism for enhancing survival during periods of illness^49^. On the other hand, mating and sexual behaviors are energy-intensive activities that can disrupt regular sleep schedules and immune functions^18^. During mating seasons, many species exhibit reduced sleep duration and altered sleep quality, as they prioritize reproductive efforts over rest^16, 17^. However, the balance between sleep and reproductive activities is crucial, as insufficient sleep can impair cognitive and physical performance, thus reducing survival prospects. The interplay of these factors highlights a complex evolutionary trade-off in which organisms must navigate the demands of immune function, reproductive success, and restorative sleep to optimize their overall fitness and survival.

Our study provides a comprehensive analysis of the interplay between sleep-regulating genes, mating status, and pathogen resistance in *Drosophila melanogaster*. We observed significant differences in fungal loads and survival rates among sleep and circadian rhythm mutants compared to wild-type CS flies. Notably, *Clk^Jrk^* mutants were the most able to repress fungal proliferation and survived the longest, whereas the *Sh^mns^*and *per^01^* mutants showed increased susceptibility. Additionally, virgin flies survived the infection longer than mated flies, which aligned with delayed fungal proliferation. Irrespective of mutant status, males also exhibited lower fungal loads and outlived females, highlighting sex-specific differences in the immune responses.

*Drosophila* exhibits a circadian rest-activity cycle, and continuous sleep deprivation results in death^22, 23, 50^. The mechanism(s) by which sleep enhances survival is not well understood, but one plausible explanation is that sickness sleep expedites organismal recovery by switching resources from waking activities to fighting off the infection and repairing cells^28^. Most studies of the impact of sleep on illness have examined the effects of sleep deprivation^24, 51^. Conversely, the impact of Ma549 infection on enhancing sleep in wild-type flies may provide an alternative approach that examines the active function of sleep, rather than the effect of a lack of sleep.

The *Sh^mns^* mutation deployed in our study knocks out the evolutionarily conserved voltage-gated potassium channel *Shaker (Sh)*^44^, and promotes sleep through GABA transmission^43^. Our data confirmed that uninfected *Sh^mns^* mutants had significantly reduced sleep compared to WT while maintaining circadian rhythms. However, infected *Sh^mns^* flies did not sustain daytime sickness sleep and succumbed faster than WT flies, supporting our overall hypothesis that sickness sleep enhances survival. However, a striking feature of our mutant study was that *per^01^* flies, like *Sh^mns^* mutants, succumbed rapidly to infection, whereas *Clk^Jrk^*flies outlived WT flies. This is even though both the mutants block sickness sleep. Similar results were obtained by stab-inoculating *Pseudomonas aeruginosa* into the same *per^01^* and *Clk^Jrk^* mutants^34, 35, 52^. This suggests that PERIOD function has a protective role in *Drosophila* against both pathogenic bacteria and fungi, while CLK undermines this role. We previously found that resistance to Ma549 in the DGRP was correlated with defense against *P. aeruginosa*, although in contrast to Kuo et al., 2010^52^, we fed the bacteria to flies to mimic natural infections^8^. Bacteria and fungi evoke the IMD and Toll defense pathways, respectively, so we postulated that the correlation was due to a generalized multipurpose defense response to multiple pathogens^8^. The conservation of the response to *per* and *Clk* by bacteria and fungi suggests that this generalized defense includes circadian regulation. Kuo et al., 2010^52^ suggested that clock mutants may differ in their effects on survival due to CLK having additional roles besides timekeeping in fighting microbial infections. In addition to promoting *per* and *tim* expression, CLK drives the rhythmic expression of approximately 150 downstream genes^53^ that regulate other physiological processes^54^. The fact that *Clk^Jrk^* mutants were the most able to repress fungal proliferation suggests that these processes are involved in immune responses. Previous studies have shown that repression of clock-controlled genes (CCGs) can enhance defense; for example, the CCG *Achilles* represses the expression of antifungal peptides^38^, whereas the antimicrobial peptide NEMURI prolongs sleep and promotes survival after bacterial infection^49^. Other factors may also contribute to immune gene activation and deleterious effects. For example, excessive or inappropriate immune activation against *Metarhizium* spp. can shorten the lifespan of insects^55^.

In conclusion, our study highlights the critical roles of sleep and circadian rhythm genes in modulating the immune responses in *Drosophila*. We demonstrated that fungal infection in WT flies increases sickness sleep; however, knocking out the pivotal *Shaker* gene with its short sleeping phenotype but normal circadian rhythms eliminate this effect. The circadian-disrupted clock gene *per^01^* has an effect similar to that of *Sh^mns^* on survival; however, *Clk^Jrk^* undercuts this role possibly because it is pleiotropic^53^. Our findings also emphasize the importance of considering sex dimorphism and mating status in host-pathogen interactions. We previously demonstrated that some mutant *Drosophila* lines with disrupted components of the Toll and Imd immune pathways lose sex dimorphism or even reverse dimorphism, with females becoming more disease resistant^21^. Mutants with disrupted sleep and circadian rhythm genes did not reproduce this, but daytime sleep in particular exhibited significant interactions with genetic lines, sex, and disease resistance. We conclude that the interplay between circadian regulation, sleep patterns, and immune function shapes the *Drosophila* resistance to fungal pathogens, along with genetic variation and physiological state.

## Materials and Methods

### Fly stocks and virgin fly collection

*per^01^*(80928), *Clk^Jrk^* (24515), *Sh^mns^* (24149), and Canton-S (64349) flies were purchased from Bloomington *Drosophila* Stock Center (BDSC), Bloomington, Indiana. *Drosophila* were cultured on cornmeal-molasses media supplemented with yeast, agar, Tegosept, and propionic acid at 25°C and ∼70% humidity under a 12-hour light:dark cycle. Virgin flies of each genotype were collected under light CO_2_ anesthesia within 8 h of eclosion and housed in a separate vial for 2–3 days to mature.

### Fungal strains and longevity bioassays

*Metarhizium anisopliae* 549 (Ma549) was acquired from the ARS Collection of Entomopathogenic Fungal Cultures (ARSEF) (Ithaca, NY). Ma549 cultures were thawed from −80°C stock vials and grown for 10 to 14 days at 27 °C on potato-dextrose-agar media plates.

Bioassays using *M. anisopliae* 549 transformed to express green fluorescent protein (Ma549-GFP) were performed according to Lu et al., 2015^9^. Briefly, fly populations were infected at the same time of day (approximately 9 pm, shortly after lights out) and survival was monitored after topical inoculation in groups of nine replicates (25-40 flies each) per sex per line across three trials. To prepare inoculum, conidia were suspended in sterile distilled water, vortexed for 2 minutes and filtered through Miracloth (22-25µm) (Andwin Scientific). Flies (two to four days old) were vortexed with spore suspensions (20 ml, 2.5×10^4^ conidia/ml) for 30 seconds, collected by filtering through Miracloth, and transferred into vials containing fly stock food without Tegosept and propionic acid. Flies were cultured at 27°C, ∼85% relative humidity. Male and female flies were housed in separate food vials to examine sexual dimorphism and to avoid an interaction between the effect of sex and age in response to infection. Less than 5% of flies vortexed with water alone (mock-infected), or conidial suspensions died within one day, with no significant differences between lines, so flies succumbing within one day post infection were deleted from the infection data. Flies were flipped into fresh vials of food every day. The number of dead flies was recorded twice per day for 10-14 days. This method was highly reproducible with a mean LT_50_ value for control *Drs*-GFP flies of 4.682 ± 0.029, N = 126^9^. The inoculum load of spores per male and female *Drosophila* was approximately 200 colony forming units.

### Fungal colonization of hosts measured by colony forming units (CFUs)

A time course bioassay of fungal growth in the hemolymph of mated and virgin CS and sleep-mutant females and males was conducted using previously described protocols^9^. Fifteen flies per sex, mating status and line at each time point across three experiments were individually homogenized with 45 μl of 0.1% Tween 80. For resistant lines, the entire 45 μl homogenate was spread onto *Metarhizium* selective medium (Rose Bengal Agar plates supplemented with oxbile, CTAB, oxytetracycline, streptomycin, penicillin, chloramphenicol, and cycloheximide)^56^. For susceptible lines, 5 μl of a 10-fold dilution of the homogenate was spread onto each plate. Colony forming units (CFUs) were counted using the ImageJ Cell Counter up to 7-days incubation at 25°C.

### Sleep assays and data analysis

Fly populations were infected at the same time of day around 9 PM, shortly after lights-off, based on reports that flies are more resistant during nighttime after infection^36^. Locomotor activity was monitored using *Drosophila* Activity Monitors (DAM2 Trikinetics, Waltham MA). Following CO_2_ anesthesia, 32 flies per sex, mating status and line per uninfected control or treatment group were individually loaded into 5 x 65 mm monitor tubes (Trikinetics, Waltham MA) containing 5% sucrose and 2% agar. To mitigate the effects of CO_2_ anesthesia, or any other potential acclimation effects, any dead flies within the first 12 hours were discarded and replaced prior to data collection. Flies were observed for seven days in DAM2 monitors synchronized to a 12 hr light: 12 hr dark (LD) cycle with lights on at 9:00 AM EDT (Zeitgeber time = 0 h) and lights off at 9:00 PM EDT (Zeitgeber time = 12 h). Sleep data was obtained from 32 flies pooled from three experiments. Locomotor activity is recorded as infrared beam crossings in one-minute bins in DAM2 monitors. *Drosophila* sleep is defined as periods of 5 min or more of inactivity^45^. Sleep and activity parameters were collected with the DAM308 system and pre-processed by DAMFileScan113 softwares (Trikinetics), and further analyzed using custom software (Insomniac3, RP Metrix, Skillman, NJ; gift of Dr. Julie Williams, University of Pennsylvania). The DAM2 output was converted to total daily, nighttime, and daytime sleep duration represented by the number of minutes when a fly is asleep in 24 or 12 hours.

### Statistical analyses on longevity and sleep parameters

Statistical analyses were performed using GraphPad Prism 10 software (GraphPad Software, Boston, MA, USA) and JMP^®^ Pro, version 17.1 (SAS Institute Inc., Cary, NC, USA).

Host survival differences were measured as LT_50_ values (lethal time in days at which 50% of the flies had died) calculated using R 4.1.0 (R Core Team 2021). To assess the survival rates of flies following Ma549 infection, we performed Kaplan-Meier survival analysis. Survival curves were generated for each group (CS, *Sh^mns^*, *per^01^*, and Clk*^Jrk^*), including variations in sex (male and female) and mating status (mated and virgin). Kaplan-Meier plots were constructed using GraphPad Prism 10, and statistical significance between survival curves was determined using the log-rank (Mantel-Cox) test and Gehan-Breslow-Wilcoxon test. Percentage survival and standard errors at all points were calculated and compared across different genotypes and conditions to assess the impact of genetic variation and infection on fly survival.

We assessed the normality of our sleep data using the D’Agostino-Pearson normality test and Shapiro–Wilk normality test, which have great power properties over a variety of different statistical distributions^57^. For data sets that passed at least one of the normality tests and had statistically similar standard deviations (SDs), we used the unpaired Student’s t-test when comparing two groups and one-way ANOVA (Tukey’s multiple comparisons) when comparing three or more groups. Otherwise, we applied Welch’s correction (Welch’s t-test) when comparing two groups and the one-way Welch’s ANOVA tests (Dunnett’s multiple comparisons) when standard deviations (SDs) were found to be significantly different (p < 0.05). For data sets that failed the normality tests, we used the Mann–Whitney U test when comparing two groups and the Kruskal-Wallis test with Dunn’s post hoc test when comparing three or more groups. We report p-values, median ± IQR (interquartile range: the difference between the 75th and 25th percentiles) from the Kruskal-Wallis test with Dunn’s post hoc multiple comparisons test, and means ± SEM from ordinary ANOVA or Welch’s test. For all tests, statistical significance was set at p < 0.05.

For the ANOVA involving four factors and generating interaction profiles, JMP Pro software was utilized.

## Acknowledgement

MN was supported by the Hatch Project Accession No. 1015969 from the USDA National Institute of Food and Agriculture (https://nifa.usda.gov/apply-grant) awarded to RJS.

## Competing Interests

The authors declare no competing financial interests.

## Availability of materials and data

All data generated or analyzed during this study are included in this published article (and its Supplementary Information files).

## Contributions

M.N. and R.J.S. conceived of and designed experiments. M.N. performed experiments. M.N. and

R.J.S. acquired, analyzed, and interpreted data. M.N. and R.J.S. wrote and edited the manuscript.

R.J.S. obtained funding and supervised experiments. Both authors reviewed and approved the manuscript.

## Corresponding authors

Correspondence to Mintong Nan or Raymond J. St. Leger.

## Notes

### Competing Interest Statement

The authors have declared no competing interest.

